# The Children’s Brain Tumor Network (CBTN) - Accelerating Research in Pediatric Central Nervous System Tumors through Collaboration and Open Science

**DOI:** 10.1101/2022.10.14.511975

**Authors:** Jena V. Lilly, Jo Lynne Rokita, Jennifer L. Mason, Tatiana Patton, Stephanie Stefankiewiz, David Higgins, Gerri Trooskin, Carina A. Larouci, Kamnaa Arya, Elizabeth Appert, Allison P. Heath, Yuankun Zhu, Miguel A. Brown, Bo Zhang, Bailey K. Farrow, Shannon Robins, Allison M. Morgan, Thinh Q. Nguyen, Elizabeth Frenkel, Kaitlin Lehmann, Emily Drake, Catherine Sullivan, Alexa Plisiewicz, Noel Coleman, Luke Patterson, Mateusz Koptyra, Zeinab Helili, Nicholas Van Kuren, Nathan Young, Meen Chul Kim, Christopher Friedman, Alex Lubneuski, Christopher Blackden, Marti Williams, Valerie Baubet, Lamiya Tauhid, Jamie Galanaugh, Katie Boucher, Heba Ijaz, Kristina A. Cole, Namrata Choudhari, Mariarita Santi, Robert W. Moulder, Jonathan Waller, Whitney Rife, Sharon J. Diskin, Marion Mateos, Donald W. Parsons, Ian F. Pollack, Stewart Goldman, Sarah Leary, Chiara Caporalini, Anna Maria Buccoliero, Mirko Scagnet, David Haussler, Derek Hanson, Ron Firestein, Jason Cain, Joanna J. Phillips, Nalin Gupta, Sabine Mueller, Gerald Grant, Michelle Monje-Deisseroth, Sonia Partap, Jeffrey P. Greenfield, Rintaro Hashizume, Amy Smith, Shida Zhu, James M. Johnston, Jason R Fangusaro, Matthew Miller, Matthew D. Wood, Sharon Gardner, Claire L. Carter, Laura M. Prolo, Jared Pisapia, Katherine Pehlivan, Andrea Franson, Toba Niazi, Josh Rubin, Mohamed Abdelbaki, David S. Ziegler, Holly B. Lindsay, Ana Guerreiro Stucklin, Nicolas Gerber, Olena M. Vaske, Carolyn Quinsey, Brian R. Rood, Javad Nazarian, Eric Raabe, Eric M. Jackson, Stacie Stapleton, Robert M. Lober, David E. Kram, Carl Koschmann, Phillip B. Storm, Rishi R. Lulla, Michael Prados, Adam C. Resnick, Angela J. Waanders

## Abstract

Pediatric brain tumors are the leading cause of cancer-related death in children in the United States and contribute a disproportionate number of potential years of life lost compared to adult cancers. Moreover, survivors frequently suffer long-term side effects, including secondary cancers. The Children’s Brain Tumor Network (CBTN) is a multi-institutional international clinical research consortium created to advance therapeutic development through the collection and rapid distribution of biospecimens and data via open-science research platforms for real-time access and use by the global research community. The CBTN’s 32 member institutions utilize a shared regulatory governance architecture at the Children’s Hospital of Philadelphia to accelerate and maximize the use of biospecimens and data. As of August 2022, CBTN has enrolled over 4,700 subjects, over 1,500 parents, and collected over 65,000 biospecimen aliquots for research. Additionally, over 80 preclinical models have been developed from collected tumors. Multi-omic data for over 1,000 tumors and germline material is currently available with data generation for > 5,000 samples underway. To our knowledge, CBTN provides the largest open-access pediatric brain tumor multi-omic dataset annotated with longitudinal clinical and outcome data, imaging, associated biospecimens, child-parent genomic pedigrees, and *in vivo* and *in vitro* preclinical models. Empowered by NIH-supported platforms such as the Kids First Data Resource and the Childhood Cancer Data Initiative, the CBTN continues to expand the resources needed for scientists to accelerate translational impact for improved outcomes and quality of life for children with brain and spinal cord tumors.

## Introduction

Brain and other central nervous system (CNS) tumors are the leading cause of cancer-related death in children and frequently result in substantial long-term morbidity and disability (1). Despite scientific advances in brain tumor biology across a large number of tumor types and histologies, improvements in long-term survival and quality of life for CNS tumors have remained elusive compared to other childhood cancers. Barriers to therapeutic advancement include lack of clinically-annotated biospecimens, insufficient preclinical models, and siloed, disconnected datasets (2).

The mission of the Children’s Brain Tumor Network (CBTN, https://cbtn.org/), launched in 2011 as the Children’s Brain Tumor Tissue Consortium (CBTTC), is to serve as a collaborative multi-institutional research consortium with a publicly-accessible biosample and data repository dedicated to the study and treatment of childhood brain tumors. CBTN seeks to address critical unmet needs for integrated, large-scale biospecimen, multi-omic, and longitudinally clinically-annotated resources. As of 2022, the CBTN comprises 32 member institutions within the United States, Italy, Switzerland, and Australia. To overcome the challenges of global, collaborative research and siloed resources, CBTN spearheaded the development of cloud-based informatics and data applications that allow researchers to access and collaboratively analyze datasets. As such, CBTN’s foundation of “Innovation through Collaboration” is being realized through its creation of a state-of-the-art biorepository, innovative analytics platforms, and real-time sharing of data and specimens. By design, CBTN initiatives build upon the success of The Cancer Genome Atlas (TCGA) and Therapeutically Applicable Research To Generate Effective Treatments (TARGET) consortia by further developing standards for the collection of specimens and comprehensive longitudinal clinical data while addressing gaps in pediatric brain tumor representation in such resources. Recently, CBTN resources have contributed to the development of cross-disease resources such as the Gabriella Miller Kids First (GMKF) Data Resource and the NCI’s Childhood Cancer Data Initiative (CCDI).

### Collaborative biobanking with the CBTN

Patients are consented by one of 32 participating sites and enrolled on a local IRB-approved protocol which includes key language to enable prospective collection of, future research on, and sharing of, de-identified surgical specimens, patient demographics, medical history, diagnoses, treatments, and clinical imaging. CHOP reviews each site’s regulatory documents prior to submission and maintains copies along with annual approvals. When possible, site-enabled workflows support the generation of cell lines, organoids, and/or patient-derived orthotopic xenografts from available tissue (**Figure 1A**).

**Figure 1.**
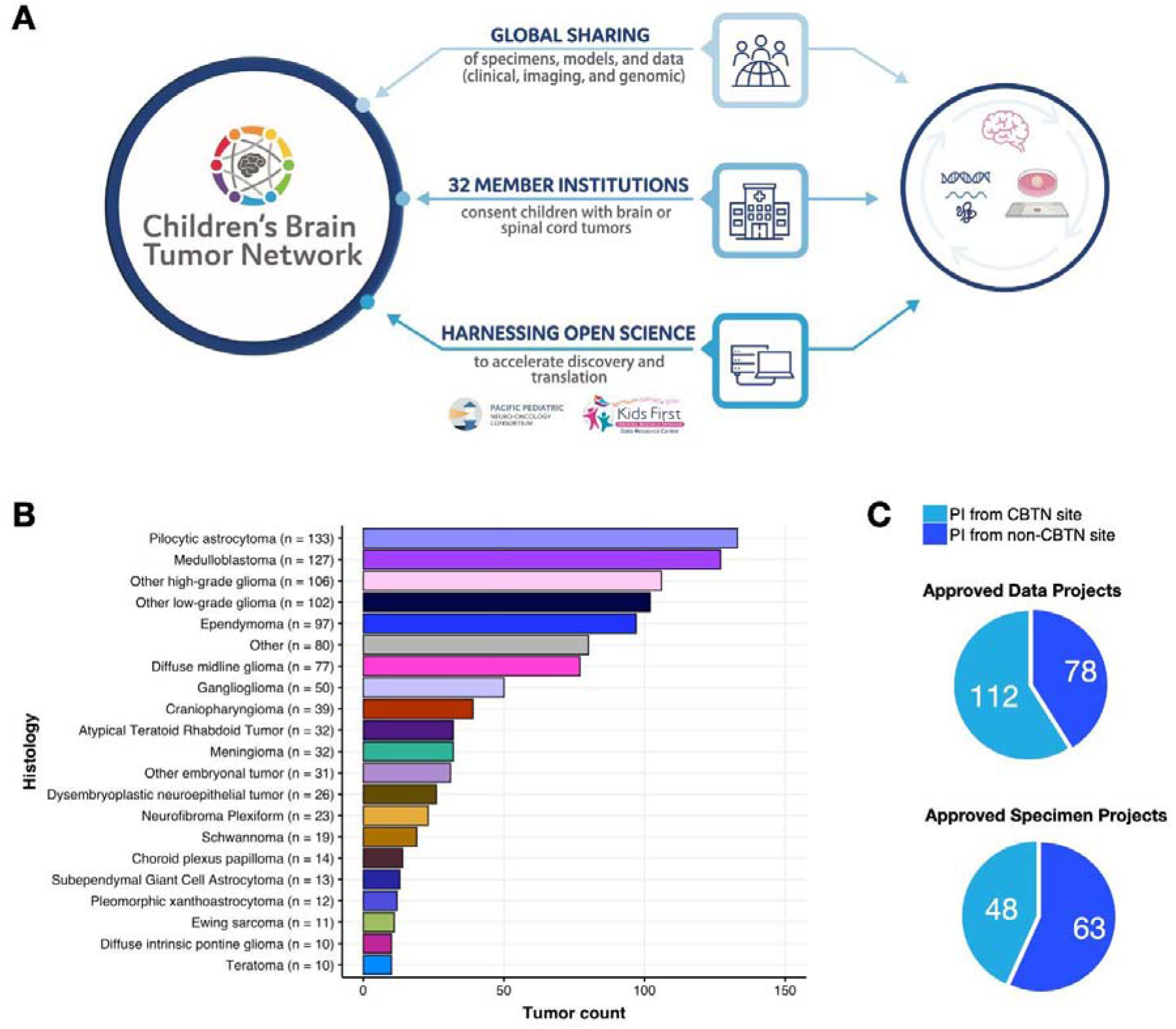
The Children’s Brain Tumor Network enables collaborative biospecimen, model, and data sharing. Shown in **A** is the CBTN ecosystem, consisting of patient enrollment at one of 32 institutions to sample and clinical data collection, biobanking, preclinical model generation, and genomic data generation to collaborative open science to accelerate discovery and clinical translation through ongoing trials such as PNOC. **B**, histologic distribution of unique tumors from the OpenPBTA (non-cancerous samples collected such as benign tumors and/or cysts are not plotted). **C-D**, number of approved data and biospecimen projects, respectively, by PI site membership. (Biorender.com used for selected images in Panel A).

In addition, CBTN’s cross-disease research platform supports the integration and management of partnered, disease-specific biospecimen and data resources including NF2 Biosolutions (https://nf2biosolutions.org/), the Embryonal Tumor with Multilayered Rosettes One Registry (https://hmh-cdi.org/etmr/), the Chordoma Foundation (https://www.chordomafoundation.org/), and OligoNation (https://www.oligonation.org/). These collaborative efforts advance disease-specific research while harnessing CBTN’s operational and research infrastructure.

### Creation of the Pediatric Brain Tumor Atlas

Created as a multi-center, multi-omic effort, the CBTN’s Pediatric Brain Tumor Atlas (PBTA) includes matched tumor-normal whole genome sequencing (WGS), tumor RNA-Seq, methylation, and proteomics, as well as longitudinal clinical data, images (MRIs, histology slides images, radiology reports), and pathology reports (3). The first PBTA dataset release of nearly 3,000 specimens from 1,074 tumors and germline sources occurred in 2018 (**Figure 1B**). A second dataset of nearly 5,000 samples including tumor/normal WGS and RNA-seq, as well as parental germline WGS, jointly sponsored by GMKF and CCDI (4), along with methylation data for > 1,700 tumors sponsored by NCI’s Center for Cancer Research, will be released with no embargo. Additionally, building upon an initial proteogenomic PBTA dataset generated in partnership with the NCI’s Clinical Proteomic Tumor Analysis Consortium (CPTAC) (5), a large > 400 sample proteogenomic cohort is underway.

In partnership with CHOP’s Center for Data Driven Discovery and Biomedicine (D3b) and the NIH GMKF Data Resource Center, the PBTA data has been integrated into cloud-based resources within the GMKF portal (http://kidsfirstdrc.org/) enabling cross-disease analysis with other GMKF datasets or those hosted by NCI’s cloud resources, such as TCGA and TARGET. In 2019, Researchers at D3b and Alex’s Lemonade Stand Foundation’s Childhood Cancer Data Lab launched the Open Pediatric Brain Tumor Atlas (OpenPBTA). OpenPBTA is a first-in-kind, open-science, collaborative analysis and manuscript-writing effort to comprehensively analyze PBTA tumors (6). OpenPBTA openly provides reproducible workflows and processed data on GitHub, PedcBioportal, and CAVATICA supporting multiple research publications as well as informing clinical trial decision-making in molecular tumor boards (7). OpenPBTA’s success has paved the way for additional efforts such as OpenPedCan (8) currently informing the new pediatric Molecular Targets Platform (https://moleculartargets.ccdi.cancer.gov/) developed with the CCDI in support of the Research to Accelerate Cures and Equity (RACE) for Children Act (9) and the Schwannomatosis Open Research Collaborative led by SAGE Bionetworks (10). CBTN promotes releasing data without embargo periods, allowing near real-time integration, dissemination, processing, and sharing of associated petabyte-scale harmonized data.

As of 2022, CBTN has supported 190 data projects and 111 biospecimen projects (**Figure 1C-D**) spanning > 25,000 biosamples. CBTN-collected biospecimens are available for request via a common approval process by both members and non-members. Such common workflows have led to the publication of >100 scientific articles and 25 abstracts in >30 peer-reviewed journals.

### CBTN partners across the globe

Embedded in the mission and vision of CBTN is the notion that collaboration is key for accelerated discoveries required to improve clinical outcomes. CBTN has benefitted from many academic, commercial, government, and advocacy partnerships (**Table 1**), empowering both molecularly-based therapeutic development and decision support as part of drug-repurposing initiatives (7) through institutional initiatives and clinical trials, such as the Pacific Pediatric Neuro-Oncology Consortium (PNOC,https://pnoc.us/).

**Table 1.**
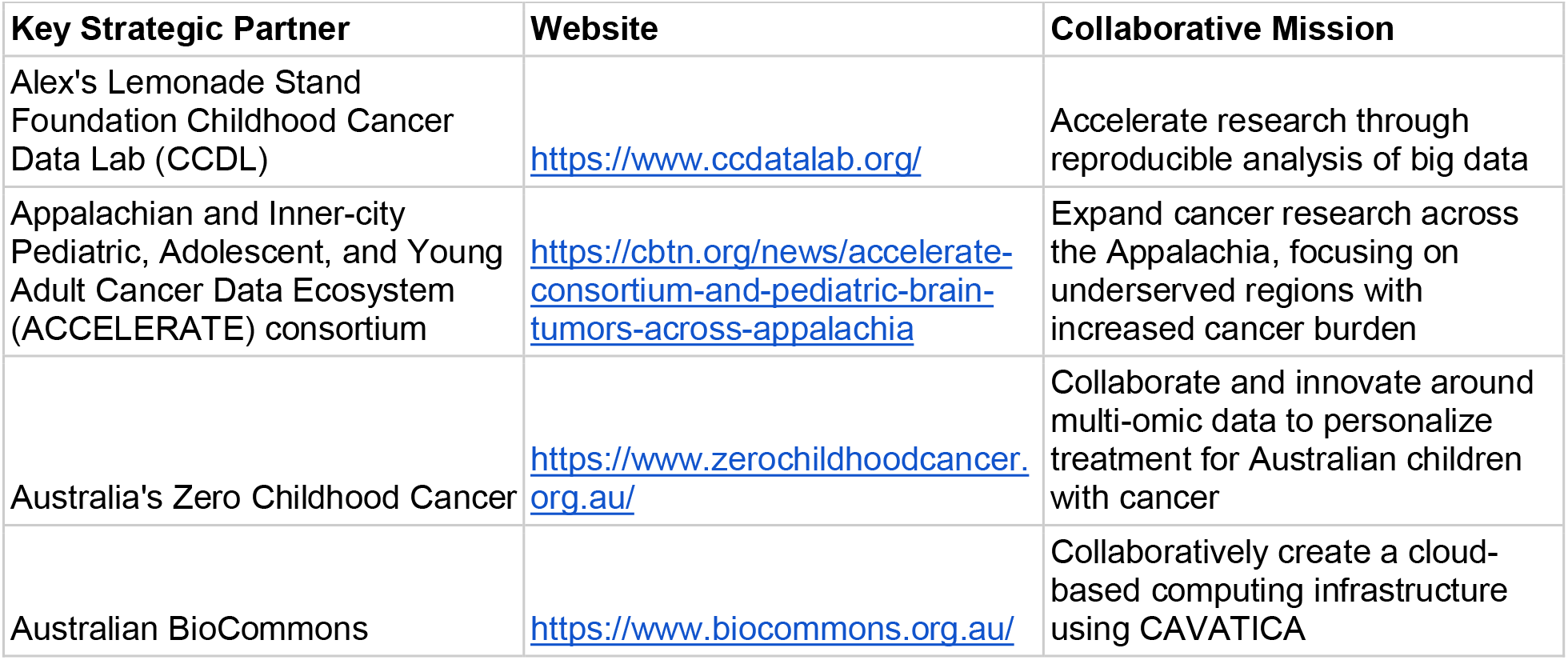
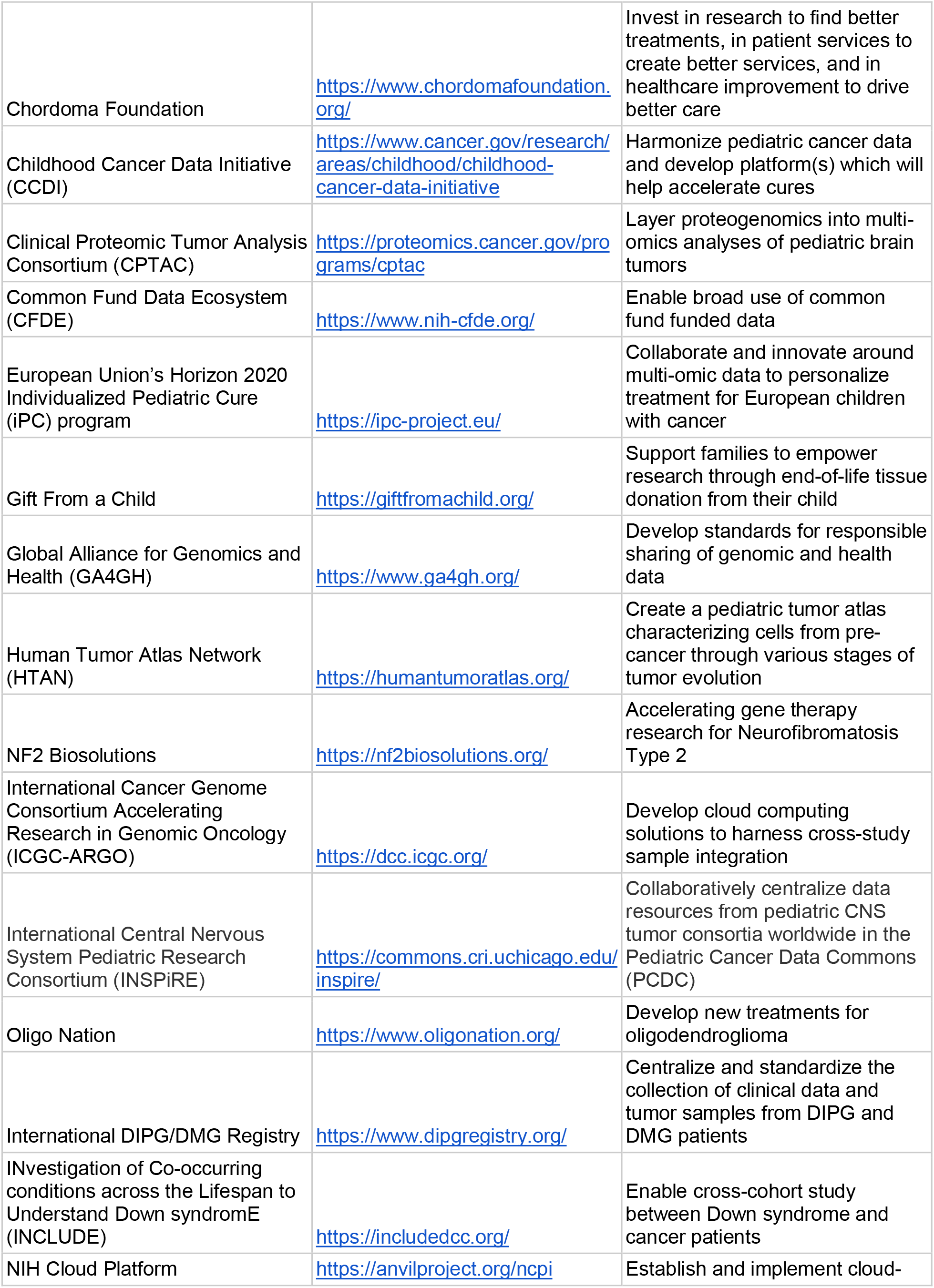
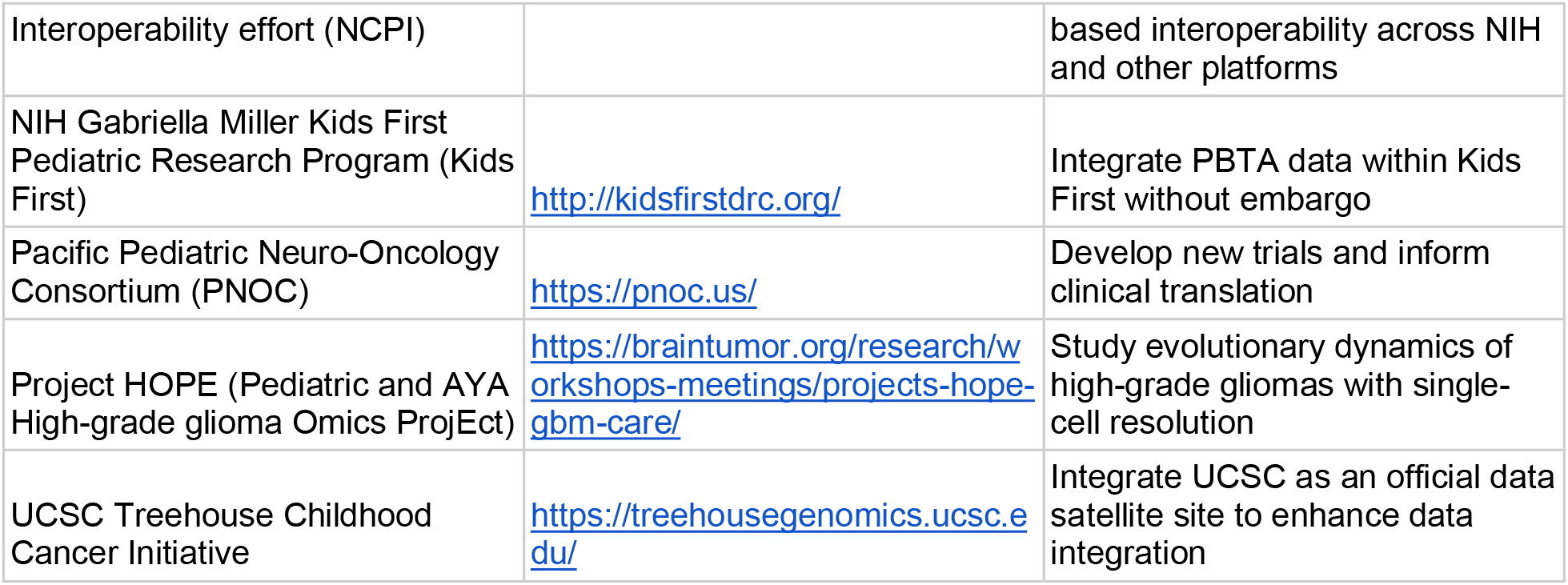
Key CBTN Strategic Partnerships. Listed below are major national and international partners, their websites, and our collaborative goals.

## Discussion

CBTN’s successes to date are empowered by its commitment to partnering with families and advocates supporting the sharing of biospecimens and data on behalf of accelerating clinical translation. CBTN, together with its partners, has developed a combination of open-science governance and platform resources that support the largest, accessible genomic and proteomic pediatric brain tumor data repository annotated with longitudinal clinical data, pathology reports and histologic images, MRI reports and images, and available biospecimens and preclinical models.

CBTN’s support of > 300 research projects generated reagents, models, data, and publications that have, in turn, enriched the CBTN’s offerings. Likewise, consortium-wide efforts towards foundational data generation like the PBTA, in combination with cloud-based platforms, support a dynamic research ecosystem that continually increases the volume and rate of brain tumor research, accelerates the development of clinical trials, and provides decision support resources to improve the outcomes of children diagnosed with CNS tumors. Importantly, CBTN is also poised to help support enforcement of the RACE Act (9), which requires companies to test cancer drugs in children that are used in adults when there is a shared molecular target. The CBTN collaborative framework of governance and paired technology advances rapid data sharing and clinical translation, defining a new model for research that breaks the traditional mold of siloed, individual achievement. Together with the many institutions, patient families, foundations, and community stakeholders, CBTN will improve the outcomes for children affected with brain cancer.

## Conflicts of interest

David S. Ziegler is a consultant, or on the advisory board, of Bayer, AstraZeneca, Accendatech, Novartis, Day One, FivePhusion, Amgen, Alexion, and Norgine. Angela J. Waanders is on the advisory board of Alexion and Day One.

## Acknowledgments

We would like to thank the patients and families participating in CBTN. CBTN is in large part philanthropically-funded, and we thank each donor for their dedication and support in making the CBTN possible. The following donors have provided leadership level support: Children’s Brain Tumor Foundation, Eaise Family Foundation, Kortney Rose Foundation, Lilabean Foundation, Minnick Family Charitable Fund, Perricelli Family, Psalm 103 Foundation, and Swifty Foundation. CBTN’s leadership would like to recognize and thank the early leadership and foundational contributions of Drs. Tom Curran, Ph.D., FRS (currently at Children’s Mercy Kansas City) and Peter Phillips, M.D. (formerly at CHOP, retired) to the creation of Children’s Brain Tumor Tissue Consortium (CBTTC) which has evolved into the Children’s Brain Tumor Network (CBTN).

## Children’s Brain Tumor Network Members

Past and present members of CBTN’s Executive Council and CHOP’s ‘s Brain Tumor Board of Visitors who inspired the creation and ensured the sustainability of CBTN are Alan Stalling, Jr., Al Gustafson, Al Musella, Amanda Haddock, Amy Summy, Amy Weinstein, Amy Wood, Andrea Gorsegner, Anita Nirenberg, Ann Friedholm, Bob Budlow, Caroline Court, Carrie Ann Stallings, Charles Genaurdi, Jr., Daniel Hare, Daniel Lipka, David Bovard, Dean Crowe, Deborah Eaise, Eliza Greenbaum, Gerald Kilhefner, Geralyn Ryerson, Ginny McLean, Graham Cox, Heather Ward, Hank Summy, James Blauvelt, James Minnick, James Ryerson, Jeannine Norris, Jessica Kilhefner, John Nilon, Kevin Eaise, Kim Hare, Kim MacNeill, Kim Wark, Kristen Gillette, Laura Cooke, Leigh Anna Lang, Lisa Ward, Liz Dawes, Mario Lichtenstein, Mark Mosier, Meghan Gleeson, Meghan Gould, Nancy Minnick, Nicole Giroux, Patti Gustafson, Patricia Genuardi, Paula Olson, Paul Touhey, Peter Norris, Richard Haddock, Robert Martin, Sarah Lilly, Scott Perricelli, Stacia Wagner, Stephanie Strotbeck, Stephanie Marvel, Stephan Ward, Sue Perricelli, Susan Funck, Timothy Court, Toni HeadTrisha Danze, W. Craig Marvel, and Wendy Payton.

Past and present members of CBTN who contributed to the generation of biospecimens and clinical and/or genomic data are Adam A. Kraya, Adam C. Resnick, Alex Felmeister, Alexa Plisiewicz, Allison M. Morgan, Allison P. Heath, Amanda Toke, Ammar S. Naqvi, Avi Kelman, Alex Felmeister, Alex Gonzalez, Alyssa Paul, Amanda Saratsis, Amy Smith, Ana Aguilar, Ana Guerreiro Stücklin, Anastasia Arynchyna, Andrea Franson, Angela J. Waanders, Angela N. Viaene, Anita Nirenberg, Anna Maria Buccoliero, Anna Yaffe, Anny Shai, Anthony Bet, Antoinette Price, Arlene Luther, Ashley Plant, Augustine Eze, Bailey K. Farrow, Baoli Hu, Beth Frenkel, Bo Zhang, Bonnie Cole, Brian M. Ennis, Brian R. Rood, Brittany Lebert, Caralyn Higginbottom, Carina A. Larouci, Carl Koschmann, Caroline Caudill, Caroline Drinkwater, Carrie Coleman-Campbell, Cassie N. Kline, Catherine Sullivan, Chanel Keoni, Chiara Caporalini, Christine Bobick-Butcher, Christopher Mason, Chunde Li, Claire L. Carter, Claudia MaduroCoronado, Clayton Wiley, Colleen Raftery, Cynthia Wong, Dan Kolbman, David E. Kram, David Haussler, David Pisapia, David Stokes, David S. Ziegler, Denise Morinigo, Derek Hanson, Donald W. Parsons, Elizabeth Appert, Emily Drake, Emily Golbeck, Emma Connell, Ena Agbodza, Eric H. Raabe, Eric M. Jackson, Erin Alexander, Esteban Uceda, Eugene Hwang, Fausto Rodriquez, Gabrielle S. Stone, Gary Kohanbash, Gavriella Silverman, George Rafidi, Gerald Grant, Gerri Trooskin, Gilad Evrony, Graham Keyes, Hagop Boyajian, Holly B. Lindsay, Holly C. Beale, Holly Sammartino, Ian F. Pollack, James Johnston, James Palmer, Jane Minturn, Jared Donahue, Jared Pisapia, Jason E. Cain, Jason R. Fangusaro, Javad Nazarian, Jeanette Haugh, Jeff Stevens, Jeffrey P. Greenfield, Jeffrey Rubens, Jena V. Lilly, Jennifer L. Mason, Jessica B. Foster, Jessica Cuba, Jessica Legaspi, Jim Olson, Jo Lynne Rokita, Joanna J. Phillips, Jonathan Waller, Josh Rubin, Judy E. Palma, Justin McCroskey, Justine Rizzo, Kaitlin Lehmann, Kamnaa Arya, Karlene Hall, Katherine Pehlivan, Ken Mosby, Kenneth Seidl, Kimberly Diamond, Komal S. Rathi, Kristen Harnett, Kristina A. Cole, Krutika S. Gaonkar, Kundan Kunapareddy, Lamiya Tauhid, Laura Prolo, Leah Holloway, Leslie Brosig, Lina Lopez, Lionel Chow, Madhuri Kambhampati, Mahdi Sarmady, Margaret Nevins, Mari Groves, Mariarita Santi-Vicini, Marilyn M. Li, Marion Mateos, Mateusz Koptyra, Matija Snuderl, Matthew Miller, Matthew Sklar, Matthew D. Wood, Meghan Connors, Melissa Williams, Meredith Egan, Michael D. Kelly, Michael Fisher, Michael Koldobskiy, Michelle Monje, Migdalia Martinez, Miguel A. Brown, Mike Prados, Mike Wilder, Miriam Bornhorst, Mirko Scagnet, Mohamed AbdelBaki, Monique Carrero-Tagle, Nadia Dahmane, Nalin Gupta, Namrata Choudhari, Natasha Singh, Nathan Young, Nicholas A. Vitanza, Nicholas Tassone, Nicholas Van Kuren, Nicolas Gerber, Nithin D. Adappa, Nitin Wadhwani, Noel Coleman, Obi Obayashi, Olena M. Vaske, Olivier Elemento, Oren Becher, Parimala Killada, Phanindra Kuncharapu, Philbert Oliveros, Phillip B. Storm, Pichai Raman, Prajwal Rajappa, Remo Williams, Rintaro Hashizume, Rishi R. Lulla, Robert Keating, Robert M. Lober, Robert (Bobby) Moulder, Ron Firestein, Sabine Mueller, Sameer Agnihotri, Samuel G. Winebrake, Samuel Rivero-Hinojosa, Sarah Diane Black, Sarah Leary, Schuyler Stoller, Shannon Robins, Sharon Gardner, Shelly Wang, Sherri Mayans, Sherry Tutson, Shida Zhu, Sofie R. Salama, Sonia Partap, Sonika Dahiya, Sriram Venneti, Stacie Stapleton, Stephani Campion, Stephanie Stefankiewicz, Stewart Goldman, Susan Jones, Swetha Thambireddy, Tatiana S. Patton, Teresa Hidalgo, Theo Nicolaides, Thinh Q. Nguyen, Thomas W. McLean, Tiffany Walker, Toba Niazi, Tobey MacDonald, Valeria Lopez-Gil, Valerie Baubet, Whitney Rife, Xiao-Nan Li, Xiaoyan Huang, Ximena P. Cuellar, Xu Zhu, Yiran Guo, Yuankun Zhu, and Zeinab Helil.

## Author Contributions

All authors listed on this manuscript made significant contributions to the creation of the Children’s Brain Tumor Network through one or more of: patient consent, collection of specimens and longitudinal clinical data, and/or generation of software and data resources for the brain tumor community.

| **Author** | **Contributions** |
| --- | --- |
| Jena V. Lilly | Writing - original draft, Conceptualization, Project administration, Supervision, Data curation, Resources |
| Jo Lynne Rokita | Writing - original draft, Formal Analysis, Visualization, Writing - review & editing, Investigation, Data curation |
| Jennifer L. Mason | Writing - original draft, Project administration, Supervision, Data curation, Resources, Investigation |
| Tatiana Patton | Writing - review & editing, Project administration, Supervision, Data curation, Resources |
| Stephanie Stefankiewiz | Project administration, Methodology, Writing - original draft |
| David Higgins | Project administration, Methodology, Resources |
| Gerri Trooskin | Project administration, Methodology, Funding acquisition |
| Carina A. Larouci | Data curation, Resources |
| Kamnaa Arya | Project administration, Writing - original draft, Resources |
| Elizabeth Appert | Data curation, Resources, Writing - original draft |
| Allison P. Heath | Software, Validation, Visualization |
| Yuankun Zhu | Software, Validation, Visualization |
| Miguel A. Brown | Software, Validation, Visualization |
| Bo Zhang | Software, Validation, Visualization |
| Bailey K. Farrow | Project administration, Validation, Visualization |
| Shannon Robins | Data curation, Resources, Investigation |
| Allison M. Morgan | Data curation, Resources, Project administration |
| Thinh Q. Nguyen | Data curation, Resources, Investigation |
| Elizabeth Frenkel | Data curation, Resources, Investigation |
| Kaitlin Lehmann | Data curation, Resources, Project administration |
| Emily Drake | Data curation, Resources, Investigation |
| Catherine Sullivan | Data curation, Resources, Supervision |
| Alexa Plisiewicz | Data curation, Resources, Project administration |
| Noel Coleman | Data curation, Resources, Project administration |
| Luke Patterson | Data curation, Resources, Project administration |
| Mateusz Koptyra | Project administration, Supervision, Resources, Methodology |
| Zeinab Helili | Data curation, Resources, Supervision |
| Nicholas Van Kuren | Data curation, Validation, Visualization |
| Nathan M. Young | Data curation, Validation, Visualization |
| Meen Chul Kim | Data curation, Software, Validation |
| Christopher Friedman | Data curation, Validation, Software |
| Alex Lubneuski | Software, Validation, Visualization |
| Christopher Blackden | Software, Validation, Visualization |
| Marti Williams | Data curation, Resources, Project administration |
| Valerie Baubet | Resources, Methodology, Project administration |
| Lamiya Tauhid | Resources, Methodology |
| Jamie Galanaugh | Data curation, Resources |
| Katie Boucher | Resources, Methodology |
| Heba Ijaz | Resources, Methodology |
| Kristina A. Cole | Resources, Methodology |
| Namrata Choudhari | Resources, Methodology |
| Mariarita Santi | Supervision, Resources, Project administration |
| Robert W. Moulder | Project administration, Resources |
| Jonathan Waller | Project administration, Resources |
| Whitney Rife | Project administration, Resources |
| Sharon J. Diskin | Formal Analysis |
| Marion Mateos | Project administration, Supervision, Data curation, Resources |
| Donald W. Parsons | Project administration, Supervision, Data curation, Resources |
| Ian F. Pollack | Project administration, Supervision, Data curation, Resources |
| Stewart Goldman | Project administration, Supervision, Data curation, Resources |
| Sarah Leary | Project administration, Supervision, Data curation, Resources |
| Chiara Caporalini | Data curation, Resources |
| Anna Maria Buccoliero | Project administration, Supervision, Data curation, Resources |
| Mirko Scagnet | Project administration, Supervision, Data curation, Resources |
| David Haussler | Project administration, Supervision, Formal Analysis |
| Derek Hanson | Project administration, Supervision, Data curation, Resources |
| Ron Firestein | Project administration, Supervision, Data curation, Resources |
| Jason Cain | Project administration, Supervision, Data curation, Resources |
| Joanna J. Phillips | Project administration, Supervision, Data curation, Resources |
| Nalin Gupta | Project administration, Supervision, Data curation, Resources, Writing - review & editing |
| Sabine Mueller | Project administration, Supervision, Data curation, Resources |
| Gerald Grant | Project administration, Supervision, Data curation, Resources |
| Michelle Monje-Deisseroth | Project administration, Supervision, Data curation, Resources |
| Sonia Partap | Project administration, Supervision, Data curation, Resources |
| Jeffrey P. Greenfield | Project administration, Supervision, Data curation, Resources |
| Rintaro Hashizume | Project administration, Supervision, Data curation, Resources |
| Amy Smith | Project administration, Supervision, Data curation, Resources |
| Shida Zhu | Project administration, Supervision, Data curation, Resources |
| James M. Johnston | Project administration, Supervision, Data curation, Resources |
| Jason R. Fangusaro | Project administration, Supervision, Data curation, Resources |
| Matthew Miller | Project administration, Supervision, Data curation, Resources |
| Matthew D. Wood | Project administration, Supervision, Data curation, Resources, Writing - review & editing |
| Sharon Gardner | Project administration, Supervision, Data curation, Resources |
| Claire L. Carter | Project administration, Supervision, Data curation, Resources |
| Laura M. Prolo | Project administration, Supervision, Data curation, Resources |
| Jared Pisapia | Project administration, Supervision, Data curation, Resources |
| Katherine Pehlivan | Project administration, Supervision, Data curation, Resources |
| Andrea Franson | Project administration, Supervision, Data curation, Resources |
| Toba Niazi | Project administration, Supervision, Data curation, Resources |
| Josh Rubin | Project administration, Supervision, Data curation, Resources |
| Mohamed Abdelbaki | Project administration, Supervision, Data curation, Resources |
| David S. Ziegler | Project administration, Supervision, Data curation, Resources |
| Holly B. Lindsay | Project administration, Supervision, Data curation, Resources |
| Ana Guerreiro Stucklin | Project administration, Supervision, Data curation, Resources |
| Nicolas Gerber | Project administration, Supervision, Data curation, Resources |
| Olena M. Vaske | Project administration, Supervision, Data curation, Resources |
| Carolyn Quinsey | Project administration, Supervision, Data curation, Resources |
| Brian R. Rood | Project administration, Supervision, Data curation, Resources, Writing - review & editing |
| Javad Nazarian | Project administration, Supervision, Data curation, Resources |
| Eric Raabe | Project administration, Supervision, Data curation, Resources |
| Eric M. Jackson | Project administration, Supervision, Data curation, Resources |
| Stacie Stapleton | Project administration, Supervision, Data curation, Resources |
| Robert M. Lober | Project administration, Supervision, Data curation, Resources |
| David E. Kram | Project administration, Supervision, Data curation, Resources |
| Carl Koschmann | Project administration, Supervision, Data curation, Resources, Writing - review & editing |
| Phillip B. Storm | Project administration, Supervision, Data curation, Resources |
| Rishi R. Lulla | Project administration, Supervision, Data curation, Resources |
| Michael Prados | Project administration, Supervision, Data curation, Resources, Writing - review & editing |
| Adam C. Resnick | Conceptualization, Project administration, Supervision, Data curation, Resources |
| Angela J. Waanders | Writing - original draft, Project administration, Supervision, Data curation, Resources, Writing - review & editing |

## Notes

### Competing Interest Statement

David S. Ziegler is a consultant, or on the advisory board, of Bayer, Astra Zeneca, Accendatech, Novartis, Day One, FivePhusion, Amgen, Alexion, and Norgine.

### Summary of Updates

- Update one author last name - Add ORCIDs, where available - Add author contributions table - Add CBTN members to back matter

